# Improved accuracy and precision of bioprinting through progressive cavity pump-controlled extrusion

**DOI:** 10.1101/2020.01.23.915868

**Authors:** Philipp Fisch, Martin Holub, Marcy Zenobi-Wong

**Affiliations:** Department of Health Science and Technology, ETH Zurich, Zurich, Switzerland; Department of Bionanoscience, TU Delft, Delft, Netherlands

**Keywords:** bioprinting, progressive cavity pump

## Abstract

3D bioprinting has seen a tremendous growth in recent years in a variety of fields such as tissue and organ models, drug testing and regenerative medicine. This growth has led researchers and manufacturers to continuously advance and develop novel bioprinting techniques and materials. Although new bioprinting methods are emerging (e.g. contactless and volumetric bioprinting), micro-extrusion bioprinting remains the most widely used method. Micro-extrusion bioprinting, however, is still largely dependent on the conventional pneumatic extrusion process, which relies heavily on homogenous biomaterial inks and bioinks to maintain a constant material flowrate. Augmenting the functionality of the bioink with the addition of nanoparticles, cells or biopolymers can induce inhomogeneities resulting in uneven material flow during printing and/or clogging of the nozzle, leading to defects in the printed construct. In this work, we evaluated a novel extrusion technique based on a miniaturized progressive cavity pump. We compared the accuracy and precision of this system to the pneumatic extrusion system and tested both for their effect on cell viability after extrusion. The progressive cavity pump achieved a significantly higher accuracy and precision compared to the pneumatic system while maintaining good viability and was able to maintain its reliability independently of the bioink composition, printing speed or nozzle size. Progressive cavity pumps are a promising tool for bioprinting and could help provide standardized and validated bioprinted constructs while leaving the researcher more freedom in the design of the bioinks with increased functionality.

## 1. Introduction

3D bioprinting with its potential to position materials and cells in a precise 3-dimensional arrangement has gained growing interest for use in tissue and organ models, drug testing and regenerative medicine, leading to a tremendous growth in the bioprinting industry. [1] Starting from the first inkjet bioprinter, a modified HP 660C printer, the technology has evolved continuously and users can nowadays choose from a variety of commercially available bioprinting technologies. [2-4]

These processes can be categorized based on the 4 main governing strategies: (i) laser induced forward transfer (ii) droplet-based bioprinting, (iii) extrusion-based bioprinting and (iv) stereolithography-based bioprinting, each of which can be further sub-categorized based on the exact mechanisms with which material and cells are positioned. [1] Among these, extrusion-based bioprinting is the most commonly used process to fabricate tissues and organs, mainly attributed to its ease of use, scalability and wide range of printable materials. The process functions by positive displacement of the material via a plunger either driven by pneumatic pressure (pneumatic extrusion) or a piston which displaces the plunger (piston driven extrusion). Another method frequently referred to in literature is the use of an Auger screw which transports material from a feedstock to the nozzle, but so far this has only been described in the context of conventional fused deposition modeling 3D printing. [3, 5-7]

Despite the ability of extrusion-based bioprinting to print a wide range of hydrogel precursors, great care in the composition and preparation of the printing materials is required to obtain the rheological and biological characteristics needed for excellent printability and biocompatibility. [8] Furthermore, additional factors such as the mixing of the bioink or the removal of air bubbles are critical, as uneven flow and nozzle clogging due to inhomogeneities in the bioink are a commonly observed phenomenon, and pneumatic extrusion systems are particularly susceptible. [1, 9] Especially highly viscous biomaterial inks and bioinks containing viscosifiers as well as micro- and nano-composite bioinks are difficult to print. Without online adjustment of the pressure to compensate for variations in the volume flow due to these inhomogeneities, the structural integrity as well as the shape fidelity of the bioprinted construct cannot be guaranteed. This becomes important for clinical applications of patient-specific bioprinting where regulatory authorities will demand standardization of the printing process. [10]

Progressive cavity pumps (PCPs) offer an alternative to the aforementioned pneumatic extrusion-bioprinting process (Figure 1 A). Initially used in heavy industry for tasks such as artificial lifting in the oil industry, mining operations or food processing, their miniaturization opened up their widespread use in fields such as adhesive and sealant applications, medical device manufacturing and 3D printing. [11-13] Recently their use in 3D printing of materials incorporating complex compositional and mechanical gradients using a two-component PCP system has been explored. [14, 15] The extrusion process of PCPs is governed by a single-helical rotor which rotates eccentrically within a double-helical stator of twice the pitch length (Figure 1 B). Between the seal lines (the contact lines of rotor and stator), cavities are created which are constantly moved towards the discharge end of the pump. As one cavity is eliminated, another cavity develops, keeping the cross-sectional area of the cavities constant, leading to a continuous non-pulsating flow. [16] The developed pushing– and–suction action therefore allows PCPs to exert significantly lower shear rates on the pumped material, compared to other pump types, such as membrane pumps or peristaltic pumps. Additionally, by knowing the geometry of rotor and stator, the flow rate and deposited volume can extremely accurately be controlled by the rotational speed of the rotor, e.g. down to 1 μl. [17] The successful application of PCPs for 3D printing of biological materials has been shown by printing bacterial spores to create biohybrid films [18, 19] and by printing bacteria in precise 3-dimensional shapes for biomedical applications. [20]

**Figure 1:**
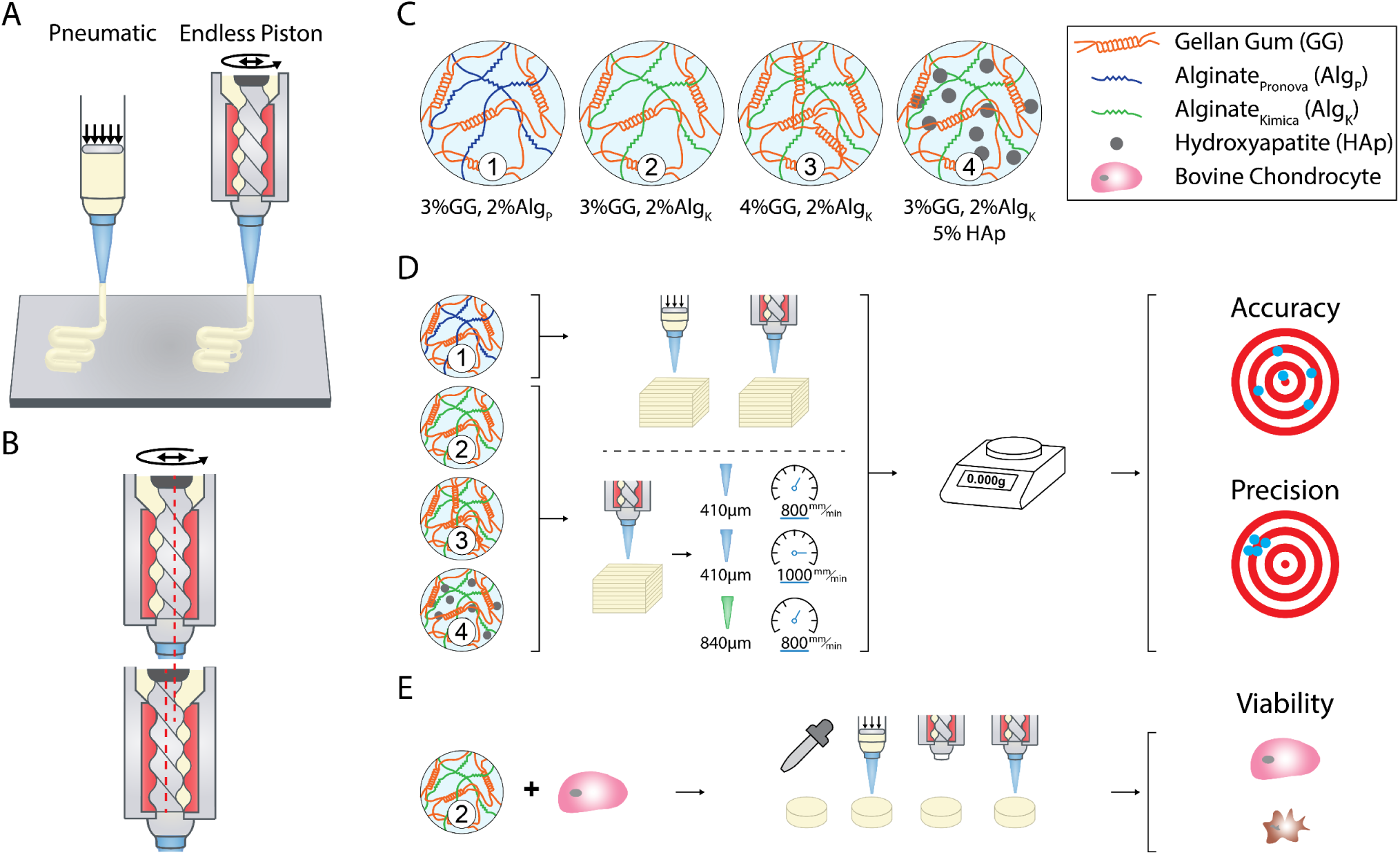
**A**, Illustration of the two different extrusion modes compared in this study. **B**, Sequential images of material being pushed through a PCP while the rotor moves eccentrically (red dotted line), where the light-yellow regions are cavities of bioink. **C**, Schematic of the different alginate/gellan gum bioinks formulations (1–4) used in this study. **D**, Schematic of the evaluation process. The bioprinting process for Bioink 1 was first optimized for both pneumatic and PCP extrusion. Afterwards, bioprinted constructs were weighed, and the accuracy and precision of the two systems compared. Bioinks 2–4 were used to evaluate the precision and accuracy of the PCP with different printing parameters: constructs were printed with a 410 µm nozzle at 800 mm/min, 410 µm nozzle at 1000 mm/min and 840 µm nozzle at 800 mm/min. **E**, Bioprinting of Bioink 2 with 3 million bovine chondrocytes/ml. Samples were casted or bioprinted with the pneumatic system with a 410 µm nozzle and the PCP with and without a 410 µm nozzle. Viability was assessed directly and one day after bioprinting.

In this paper we compare the most commonly used pneumatic extrusion process to a miniaturized progressive cavity pump in terms of their accuracy and precision as well as the compatibility of the extrusion process with bioprinting, i.e. living cells (Figure 1A). To do so, an established bioink made from alginate and gellan gum was used. [21, 22] We further tested the sensitivity of the extrusion process to increased polymer content, addition of hydroxyapatite particles, and commercial source of alginate (Figure 1 C, Bioink 1–4). Accuracy and precision of the two methods were determined by comparing the actual weight of the printed constructs compared to the targeted weight, after both printing processes were optimized. To further test the boundaries of the PCP extrusion, we studied if the accuracy and precision could be maintained when increasing nozzle diameter and printing speed (Figure 1D). Lastly, to evaluate the effect on cell viability, constructs were casted and bioprinted with the pneumatic system or the PCP, with and without a printing nozzle, and cell viability analyzed directly and one day after printing (Figure 1 E).

## 2. Methods

### 2.1 Bioprinting

Bioprinting was performed on a Biofactory bioprinter (regenHU, Switzerland). Printing files were either prepared in slic3r (https://slic3r.org/) and post-processed in a custom written Matlab postprocessor (Matlab 2018a, Mathworks) or prepared in BioCAD (regenHU, Switzerland).

### 2.2 Progressive cavity pump (PCP) installation and preparation

The eco-PEN300 PCP (preeflow by ViscoTec GmbH) was installed in the Biofactory with a custom 3D printed mount (Supplementary Figure 1 D–F). The eco-CONTROL EC200-K unit controlled the PCP, allowing precise calibration and setting of the volume flow of the eco-PEN300. As start/stop signal, the 24 V pressure on/off signal of the DD-135N Biofactory printhead was connected to the eco-CONTROL unit.

To prepare the PCP for printing, the bioink cartridge was connected to the PCP via a Luer lock adapter allowing material flow into the cavity ahead of the rotor/stator of the PCP. Material flow into the PCP was realized by pneumatic pressure. The cartridge was connected to a pressure regulator of the bioprinter, therefore pushing material at a continuous pressure into the PCP. To vent the cavity, the bleed screw of the PCP was opened, pressure applied to the plunger of the bioink cartridge and material allowed to flow out until free of air bubbles. Afterwards, the bleed screw was closed and the PCP was started until material began to flow out of the nozzle. Before each test, the PCP was calibrated by extruding a specific amount of material, weighing it, converting the measured weight to volume and comparing the extruded volume to the expected volume.

Factors such as weighing errors, material remaining at the nozzle tip or evaporation of water from the material can influence the calibration procedure. Additionally as the calibration procedure was performed only with one volume flow (volume flow: 25% of the max. volume flow = 370 μl/min) additional test were carried out to confirm the calibration of the PCP. An amount of 50 mg was set in the control unit of the PCP and extruded at 100 μl/min, 150 μl/min and 200 μl/min (the minimum volume flow in the technical specifications of the eco-PEN300 PCP is 120 μl/min). Achievable volume flows of the pneumatic system for comparison are: 76.4 ± 9.9 μl/min (20 kPa), 129.1 ± 7.5 μl/min (25 kPa), and 212.8 ± 14.9 μl/min (30 kPa). The extrudate was collected in test tubes and weighed.

### 2.3 Bioink preparation

Gellan gum and alginate bioinks were prepared by dissolving gellan gum (Kelcogel GG-LA [GG], CP Kelco) and alginate (Algin I-1, Kimica [Alg_K_] or Pronova UP LVG [Alg_P_], Novamatrix) in 220mM d-glucose and 20mM HEPES, pH 7.4, at 90°C for 1h while stirring. Similarly, the gellan gum, alginate and hydroxyapatite particles (HAp, Acros Organics) bioink was prepared by dispersing first HAp in the aforementioned buffer after which gellan gum and alginate were added. The solution was then allowed to dissolve with stirring at 90°C for 1h. Afterwards, the solution was transferred into a beaker and continuously mixed with a spatula until cooled to RT to avoid the formation of a gel-block due to the thermal gelation of gellan gum. The obtained paste was loaded into a double syringe (L-system, medmix) and mixed 10:1 with media (DMEM 31966, gibco) through a static mixer (MLX 2.5-16-LLM, medmix), filled into a printer cartridge and centrifuged at 1500 rcf for 4 min to remove any remaining air bubbles. Four different bioinks were used. Bioink 1: 3% GG, 2% Alg_P_, Bioink 2: 3% GG, 2% Alg_K_, Bioink 3: 4% GG, 2% Alg_K_, Bioink 4: 3% GG, 2% Alg_K_, 5% HAp (Figure 1 C).

Preparation of cell containing bioinks was performed by suspending cells in media and mixing the obtained cell suspension 10:1 with the gellan gum-alginate paste to reach a final concentration of 3% GG, 2% Alg_K_ and 3 mio bovine chondrocytes per ml.

### 2.4 Rheology

Rheological characterization of the bioinks was performed on an Anton Paar MCR 310 rheometer equipped with a peltier element and thermal hood (H-PTD 200; Anton Paar, Switzerland). Before each measurement, the respective geometries were coated with poly-L-lysine (PLL) to prevent slippage between the material and the geometry. A 10 μg/ml PLL solution was placed on the plate of the rheometer and the geometry brought in close contact with the solution so that the solution was fully covering the area of the geometry. The temperature was raised to 37°C for 30 min after which the solution was removed and the bioink added.

Shear thinning behavior was characterized by performing rotational tests with a 20 mm parallel plate geometry (PP20, Anton Paar) at a gap of 0.8 mm. The shear rate was increased logarithmically from 0.01 s^-1^ to 300 s^-1^.

Shear recovery behavior was characterized by performing oscillatory tests with the same setup as in shear thinning tests. The test was split into several intervals with alternating shear strain, constant frequency of 1 Hz and measurement point duration of 10 s. In the first interval, a strain of 1% was applied for 300 s, i.e. 30 measurement points of 10 s each, after which an interval of high shear strain followed at 500% for 60 s. A short resting phase of 5 s followed to allow the material to come to rest after which the recovery of the material was measured at 1% shear strain for 900 s. The high shear strain, 5 s rest and recovery interval were then repeated once.

Stress amplitude sweep tests were performed using oscillatory tests with the same setup as in shear thinning tests. Shear stress was increased logarithmically from 0.5 Pa to 25 Pa at a frequency of 1 Hz. Yield points were calculated based on the “Yield Stress II” analysis in the Rheoplus software (Anton Paar). A tolerated deviation of 5% from the plateau stress value of the linear viscoelastic range was set as the threshold to determine the yield point. Crosslinking was characterized by performing oscillatory measurements utilizing a 10 mm parallel plate geometry (PP10, Anton Paar) at a gap of 0.8 mm. Tests were performed at 0.1% shear strain, 1 Hz and a measurement point duration of 20 s. After 5 min, 1 ml of 20 mM CaCl_2_ was added around the geometry to allow diffusion of calcium ions into the bioink and initiate crosslinking.

### 2.5 Preparation process of the pneumatic system and the progressive cavity pump

To reliably compare the two printing systems, several key parameters influencing the material deposition during bioprinting, were analyzed. These parameters included 1) die swell behavior after extrusion with the pneumatic system and the PCP, 2) volume flow rate of the bioink at different pressures for the pneumatic system, 3) extrusion delay of the individual systems before material extrusion and 4) layer height of up to 3 consecutive layers for both systems. All tests were performed with Bioink 1, where three batches of bioink were prepared for each test and each test repeated three times. Imaging was performed on a stereomicroscope (Leica Wild M650) equipped with a color camera (Leica EC3) and all images analyzed in ImageJ. To relate the extruded weight to the extruded volume, density was approximated by extruding 200 μl of bioink with a positive displacement pipette (microman, Gilson) into a test tube and weighting the material. All printing related to the comparison between pneumatic system and PCP was performed with a 410 μm nozzle.

#### 1) Die swell

as viscoelastic fluids tend to show an increase in cross-section of the extruded strand (D_ex_) compared to the cross-section of the die or nozzle which they were extruded through (D_0_)[23], the strand diameter after extrusion from a 410 μm nozzle was analyzed. This behavior can potentially influence subsequent measurements as an increase in diameter of the extrudate could lead to an increase in layer height and/or width. To measure the strand diameter after extrusion, the cartridge connected to the pneumatic system and the PCP were mounted on a stand allowing the free vertical extrusion of the bioink. Images of the extruded strand during flow were then taken at the nozzle tip and analyzed.

#### 2) Volume flow of the pneumatic system

The volume flow of the PCP can be precisely controlled by controlling the rotational speed of the rotor. [17] Contrary, the volume flow for any given pressure of the pneumatic system needs to be determined. Although approximations using mathematical models to predict the volume flow exist, these are based on preceding characterization of the shear thinning behavior of the bioink. [24] Therefore to provide a simple but fast method to measure the volume flow of the pneumatic system at any given pressure, we connected the printing cartridge to the bioprinter and extruded the bioink at a specific pressure for 30 s into test tubes after which the extruded amount was determined and the volume flow calculated.

#### 3) Extrusion delay

A delay between the start command of the gcode and the actual extrusion of material can occur in the pneumatic system due to the time it takes to open the pressure valve and the subsequent pressure buildup, and in the PCP due to the signaling delay between bioprinter, control unit of the PCP and PCP itself as well as the start of the rotor of the PCP. Before this delay could be determined, a line height and width assessment had to be performed to tune the distance of the nozzle to the glass slide to provide direct contact of the material to the glass slide and reduce the error due to material flow after extrusion. When bioinks are extruded through a nozzle, they undergo structural decomposition and recovery during the process due to shearing (shear recovery). Both behaviors are time dependent and as the bioink requires a specific time to recover its structural strength, the material tends to flow immediately after the extrusion process. This leads to a decrease in line height and increase in line width. To assess line height and width, individual lines were printed at different pressures with the pneumatic system and different volume flow rates with the PCP onto glass slides and imaged from the top and side. Line height and width were then analyzed in ImageJ. The extrusion delay was then assessed by creating a 15 mm long line in BioCAD and a delay of 0, 100, 200, 300 and 400 ms incorporated via the G4 function (interrupt program execution) right after the extrusion command. Lines were printed with different delays after which the printed line was imaged and its length analyzed in ImageJ. The optimal delay was then calculated based on the delay at which the printed line achieved the length of the line designed in BioCAD. The determined delay was used for printing consecutive layers on top of each other and for printing cubes for precision and accuracy analysis.

#### 4) Layer height

Bioprinting layers of material is not only dependent on the shear recovery behavior of the bioink but also on the fusion of strands deposited next to each other. Due to the flow immediately after extrusion, deposited strands show a larger line width than the nozzle diameter. When setting the distance between adjacent lines to the nozzle diameter, these lines therefore overlap and fuse together eventually increasing the layer height compared to the height of individual lines. To assess the height of the resulting layers, 10 lines were printed next to each other, spacing 410 μm apart. The ideal volume flow for printing these layers was determined by assuming that the fusion of strands would significantly reduce the widening of printed constructs compared to the widening of individually printed lines. Based on this assumption, one can either chose to set the volume flow and calculate the height of the construct, as the distance between adjacent lines as well as the feed rate are defined in the gcode, or one can chose the height of the construct and thereof calculate the necessary volume flow. Here, the volume flow was determined by choosing the height, assuming that the optimal print consists of strands of square cross section with side length of the square equal to the nozzle diameter. The assumption of a square cross-section stems from the immediate flow of the bioink after extrusion. As strands fuse, any cavities are filled and surface tension evens the surface of the printed construct. Using this assumption, the volume flow can be determined as follows:

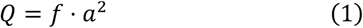

with *f* being the feed rate (mm/min) and *a* being the diameter of the printing nozzle. Applying this equation to a 410 μm nozzle and a feed rate of 800 mm/min the volume flow was determined to be 134.5 μl/min. Regarding the PCP, this volume flow can be precisely set in the control unit whereas for the pneumatic system, the pressure to achieve this volume flow (25.33 ± 0.37 kPa) was determined as described in *2)*. As the pressure sensors installed in the Biofactory had a resolution of 1 kPa and the screw to adjust the pressure did not allow a more precise adjustment, a pressure of 26 kPa was used to print with the pressure system which would lead to a volume flow of 145.8±9.0 μl/min. Due to the unknown layer height, a layer height of 410 μm was set to avoid collision of the printing nozzle with the extruded material. Up to three consecutive layers were printed on top of each other and printed constructs analyzed by imaging sideways and assessing the layer height in ImageJ.

### 2.6 Comparison of the accuracy and precision of the pneumatic system to the progressive cavity pump

Data from the preparation process were then used to design and print a test cube to compare the accuracy and precision of the pneumatic system to the PCP. The cube designed in BioCAD consisted of 26 lines spaced 410 μm apart at a length of 10.25 mm which were connected alternately on either side, therefore creating one continuous printing path per layer. The designed layer was then stacked 14 times with a distance between each consecutive layer of 410 μm. The printing path was alternately rotated by 90° on each consecutive layer. Cubes were printed at 800 mm/min (Supplementary Figure 4 A). A printing delay after the start command for extrusion of 155 ms for the pneumatic system and 284 ms for the PCP was included according to the determined extrusion delay. Additionally, a layer height of 380 μm and 403 μm was used for the pneumatic system and the PCP respectively. As described previously when determining the printing pressure for the layer height analysis, a pressure of 26 kPa was used for the pneumatic system to achieve the corresponding volume flow. After printing, the weight of each cube was immediately measured. For all tests, 3 batches of Bioink 1 were used, and 3 cubes printed with each batch and each system.

As 3D printers and bioprinters underlay the laws of physics, their print-time is influenced by the acceleration and jerk (rate of change of acceleration) of the system. [25] Therefore it is not possible to simply determine the print-time based on the length of the printing path and the feed rate, i.e. the set velocity at which the printing nozzle moves, without knowing the specification of the system. This means for example that a 10 mm line printed with a feed rate of 10 mm/s does not require 1 s to be printed but as the printer needs to accelerate to achieve the feed rate (print velocity) and decelerate to come to a stop, the print takes longer than 1 s. Additionally, at corners, the printer needs to decelerate e.g. in x-direction and accelerate in y-direction, meaning that the movement comes to a stop at these corners which can lead to increased material deposition at these positions. [25] To determine the exact volume extruded during the print process, the print-time, i.e. the time between the start and stop signal of the gcode was measured and the targeted volume calculated according to:

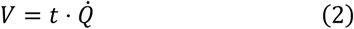

with V being the targeted volume, t being the time the pressure valve is open and 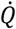 being the set volume flow. For the cubes described in this section, the print time was 333.9 s and with a volume flow of 134.5 μl/min, the targeted volume was calculated to be 748.5 μl for the PCP and with a volume flow of 145.8 μl/min, the targeted volume was 811.4 μl for the pneumatic system.

Accuracy was determined based on the percentage difference between the mean of the measured extruded volume and the targeted volume and precision determined based on the standard error.

### 2.7 Influence of the bioink composition, printing speed and nozzle diameter on the accuracy and precision of the progressive cavity pump

To compare the influence of different printing parameters and different bioinks on the precision and accuracy of the PCP, cubes were printed with different nozzle sizes and different feed rates and each test carried out with three different bioinks, Bioink 2–4 (Figure 1 D). Cubes as described in Supplementary Figure 1 A, designed for a 410 μm nozzle were printed at 800 mm/min and 1000 mm/min. Additionally, cubes designed for an 840 μm nozzle were printed at 800 mm/min. These cubes consisted of 14 lines spaced 840 μm apart at a length of 10.92 mm in a similar arrangement as in the cubes printed with a 410 μm nozzle. A total of 8 layers were printed (Supplementary Figure 4 B). All cubes were printed three times with each bioink batch and three batches of bioink were prepared for each test. All tests were carried out by calculating the desired volume flow according to Equation (1), leading to a volume flow of 134.5 μl/min for cubes printed with a 410 μm nozzle at 800 mm/min, 168.1 μl/min for cubes printed with a 410 μm nozzle at 1000 mm/min and 564.5 μl/min for cubes printed with a 840 μm nozzle at 800 mm/min. According to Equation (2) the expected volume for each cube was therefore: 748.49 μl for cubes printed with a 410 μm nozzle at 800 mm/min, 763.48 μl for cubes printed with a 410 μm nozzle at 1000 mm/min and 1034.63 μl for cubes printed with a 840 μm nozzle at 800 mm/min.

These tests were carried out without performing the printing preparation as described in *Section 2.6* to additionally test if it is sufficient to use Equation (1), provided that the bioink used is well characterized and its printability confirmed. A delay of 284 ms was used for all tests as it was assumed that the delay arises from the signaling delay between bioprinter, control unit of the PCP and PCP itself as well as the start of the rotor of the PCP and not due to nozzle size or material used. The layer height was set to the nozzle diameter as Equation (1) ideally assumes a printed line with square cross-sectional area of side length of the nozzle diameter.

### 2.7 Viability

To test the effect of the different printing modes on cell viability, Bioink 2 was combined with passage 3 bovine chondrocytes to achieve a final cell concentration of three million per milliliter. Samples were either casted or printed with the pneumatic system or the PCP with a 410 μm nozzle. To evaluate whether the stator/rotor of the PCP would damage cells or the extrusion through the nozzle, additional samples were prepared with bioink only extruded through the stator/rotor and bioink extruded through the stator/rotor and a nozzle (Figure 1 E).

Viability was assessed immediately after printing and one day after printing. Samples were washed three times in media (DMEM 31966, gibco), stained in 1uM CalceinAM and 1uM Propidium Iodide in media (DMEM 31966, gibco) for 1h and washed again three times in media (DMEM 31966, gibco). Samples were imaged on a structured illumination microscope (Zeiss Axio Observer equipped with an Apotome) from the surface of the sample 100μm into the sample with images being acquired every 5μm. Stacks were projected onto the z-plane and viability calculated by dividing the number of viable cells by the number of total cells.

### 2.8 Statistical Analysis

All statistical analysis was carried out in Matlab (Matlab 2018a, Mathworks). One- and two-way analysis of variance (ANOVA) was performed followed by multicomparison test according to the Bonferroni method. A p-value below 0.05 (p<0.05) was considered statistically significant.

## 3. Results

### 3.1 PCP installation

A confirmation of the ability of the PCP to deposit a specific volume at different volume flows was performed, as the calibration of the PCP was carried out at a higher volume flow (370 μl/min) than used in this study and as some of the measurements performed in the preparation process were conducted below the minimum volume flow of the PCP (120 μl/min) as specified in its technical specifications. For a desired 50 mg of material, the PCP extruded 49.84 ± 0.27 mg at 100 μl/min, 49.88 ± 0.30 mg at 150 μl/min and 49.92 ± 0.13 mg at 200 μl/min, which did not show a significant dependency on the volume flow (Supplementary Figure 3 D, p = 1).

### 3.2 Rheological characterization

All bioinks used in this study showed the characteristic shear recovery and shear thinning behavior with yield point necessary to achieve good printability (Supplementary Figure 2 A-L). To determine if the various bioinks differed in their rheological behavior, which ultimately determines the bioinks’ printing pressure and layer-by-layer deposition, key rheological parameters were compared. The standard deviation of these parameters provides additional information on the homogeneity of the different bioink batches and the reproducibility of the bioink preparation. Therefore, viscosity was analyzed at the initial shear rate of 0.01 s^-1^ (η_0.01_), yield stresses (τ_y_) of the different bioinks compared and the shear recovery behavior evaluated based on the storage modulus (G’) recovery and percentage recovery of G’.

A significant increase in viscosity was observed when switching form Alg_P_ to Alg_K_ (p = 0.0094). When increasing the GG concentration from 3% to 4%, the additional polymer content led to an increase in viscosity (p < 0.001). Similarly, when incorporating hydroxyapatite microparticles into the bioink, additional frictional forces occur, leading to an increase in viscosity (p < 0.001, Table 1, Supplementary Figure 2 N).

**Table 1:**
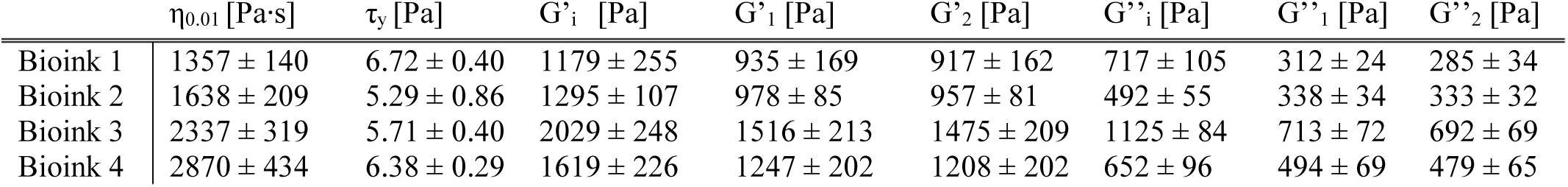
Key rheological parameters determined for the various bioinks. Viscosity at the initial shear rate of 0.01 s^-1^ (η_0.01_), yield stress (τ_y_), initial sorage modulus (G’_i_) and storage modulus after the first (G’_1_) and second (G’_2_) shear event and initial loss modulus (G’’_i_) and loss modulus after the first (G’’_1_) and second (G’’_2_) shear event.

| | $\eta_{0.01}$ [Pa·s] | $\tau_y$ [Pa] | $G'_i$ [Pa] | $G'_1$ [Pa] | $G'_2$ [Pa] | $G''_i$ [Pa] | $G''_1$ [Pa] | $G''_2$ [Pa] |
| --- | --- | --- | --- | --- | --- | --- | --- | --- |
| Bioink 1 | $1357 \pm 140$ | $6.72 \pm 0.40$ | $1179 \pm 255$ | $935 \pm 169$ | $917 \pm 162$ | $717 \pm 105$ | $312 \pm 24$ | $285 \pm 34$ |
| Bioink 2 | $1638 \pm 209$ | $5.29 \pm 0.86$ | $1295 \pm 107$ | $978 \pm 85$ | $957 \pm 81$ | $492 \pm 55$ | $338 \pm 34$ | $333 \pm 32$ |
| Bioink 3 | $2337 \pm 319$ | $5.71 \pm 0.40$ | $2029 \pm 248$ | $1516 \pm 213$ | $1475 \pm 209$ | $1125 \pm 84$ | $713 \pm 72$ | $692 \pm 69$ |
| Bioink 4 | $2870 \pm 434$ | $6.38 \pm 0.29$ | $1619 \pm 226$ | $1247 \pm 202$ | $1208 \pm 202$ | $652 \pm 96$ | $494 \pm 69$ | $479 \pm 65$ |

**Figure 2:**
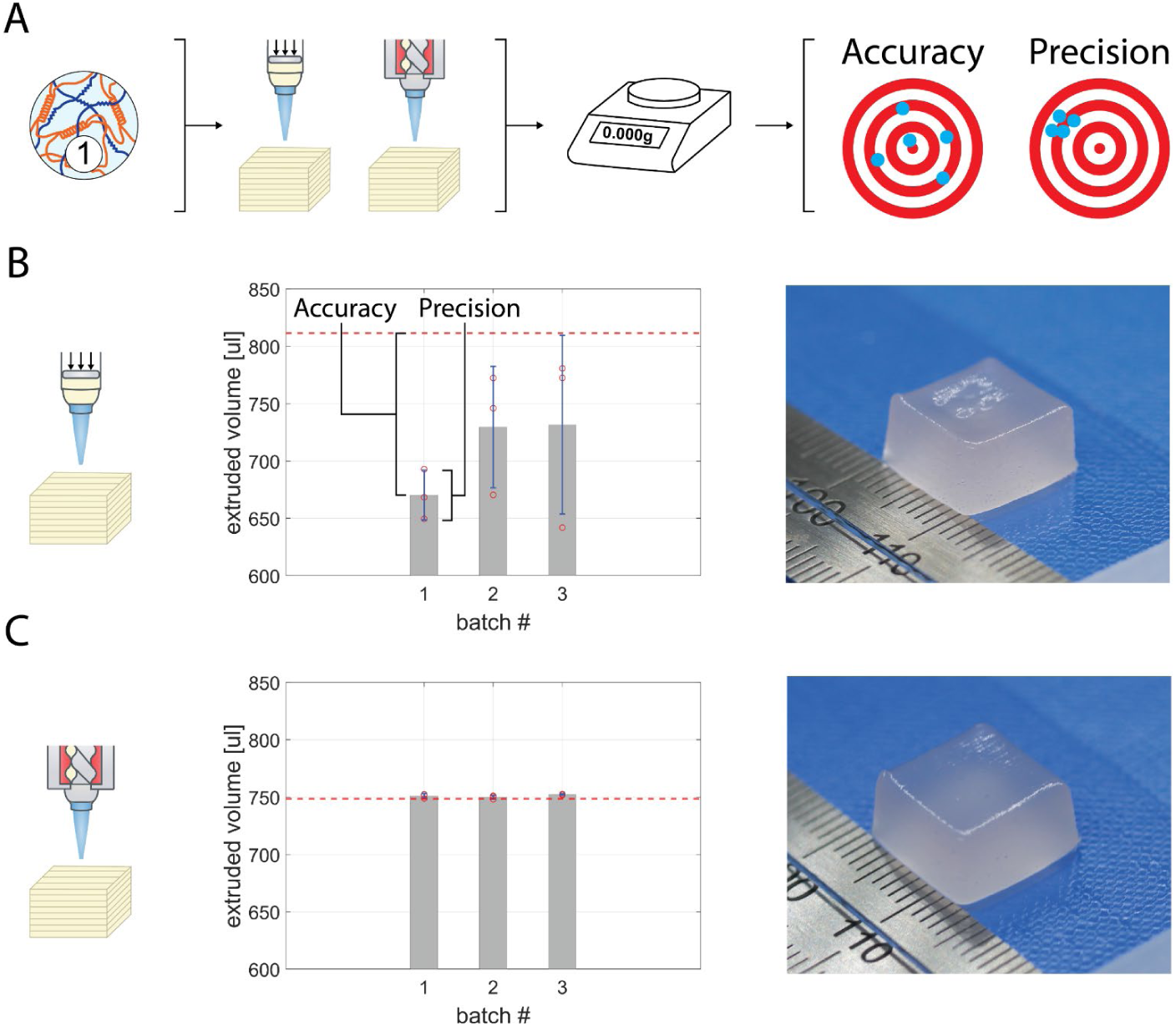
**A**, Schematic of the process followed for the comparison of the pneumatic system to the PCP. Bioink 1 was used to perform the printing assessment after which the obtained printing parameter were used to print constructs with both systems. These constructs were weighted and analysed for their precision and accuracy. **B**, Extruded volume of constructs printed with the pneumatic system. The picture shows a cube printed with the pneumatic system. A small defect can be seen on the surface on top of the cube. **C**, Extruded volume of constructs printed with the PCP. No defects were observed in the printed cubes. Red line: targeted volume.

Yield stress was higher for the bioink prepared with Alg_P_ compared to the one prepared with Alg_K_ (p < 0.001). Contrary to the viscosity analysis, no significant difference was observed in the yield stress when increasing the GG concentration from 3% to 4% (p = 0.43), whilst the addition of HAp increased the yield stress (p < 0.001, Table 1, Supplementary Figure 2 O). Analysis of the shear recovery behaviour of the bioinks was carried out based on their storage moduli (G’), which in combination with the loss modulus (G’’) provides additional information on the differences in the viscoelastic behavior of the bioinks. No significant difference in the initial G’ (index i) as well as the G’ after the first (index 1) and second (index 2) shear event occurred when changing the alginate origin from Alg_P_ to Alg_K_ (p_i_ = 0.74, p_1_ = 1, p_2_ = 1) whereas Bioink 2 showed a significantly lower G’’ before shearing but not after the first and second shear event (p_i_ = 0.01, p_1_ = 0.61, p_2_ = 0.12). Increasing the GG concentration significantly increased all storage moduli (p_i_ = 1.3e-8, p_1_ = 2.9e-7, p_2_ = 4.6e-7) as well as loss moduli (p_i_ = 4.9e-15, p_1_ = 2.4e-14, p_2_ = 4.1e-14). Similarly, the addition of HAp to Bioink 2 increased both G’ (p_i_ = 0.006, p_1_ = 0.008, p_2_ = 0.012) and G’’ (p_i_ = 3.1e-4, p_1_ = 1.2e-5, p_2_ = 2.5e-5) significantly but did not increase it as strong as the increase in GG concentration (Table 1, Supplementary Figure 2 P). Despite these differences, no significant differences were found in the percentage shear recovery of G’ after the first shear event (Bioink 1: 79.7 ± 4.1%, Bioink 2: 75.5 ± 1.0%, Bioink 3: 74.6 ± 2.2%, Bioink 4: 76.8 ± 2.2%) and only a significant difference between Bioink 1 and 3 (p = 0.025) was found after the second shear event (Bioink 1: 78.3 ± 4.4%, Bioink 2: 73.9 ± 0.9%, Bioink 3: 72.5 ± 2.1%, Bioink 4: 74.4 ± 2.4%).

### 3.3 Preparation process of the pneumatic system and the progressive cavity pump

#### 1) Die swell

Measurements of the strand diameter after extrusion through a 410 μm nozzle with the pneumatic system did not show any significant dependence on the extrusion pressure except for material extruded at 15 kPa (d = 428 ± 10 μm). A significantly smaller diameter compared to material extruded at 25 kPa (d = 439 ± 9 μm, p = 0.04) and at 30 kPa (d = 441 ± 7 μm, p = 0.01) was observed (Supplementary Figure 3 A). No significant differences were found in strand diameter when extruded with the PCP between different volume flows (Supplementary Figure 3 B, p > 0.3). The overall diameter of bioink extruded with the pressure system was 6% (436 ± 9 μm) and with the PCP 5% (431 ± 6 μm) larger than the nozzle diameter. As this increase in diameter can possibly be explained by velocity profile rearrangements and/or mass balance considerations [23], die swell was neglected in the further preparation process.

#### 2) Volume flow of the pneumatic system

Increasing the pressure at which bioink is extruded significantly increased the volume flow (Supplementary Figure 3 C). A pressure of 25.33 ± 0.37 kPa was determined to achieve a volume flow of 134.5 μl/min, corresponding to an optimal print with a 410 μm nozzle and a feed rate of 800 mm/min. Due to the resolution of the pressure sensor of the Biofactory (1 kPa) and the imprecision of the screw to adjust the pressure, the pressure was rounded up to 26 kPa which would lead to a volume flow of 145.8 μl/min.

#### 3) Extrusion Delay

Line height and line width analysis were performed to optimize the distance of the nozzle to the glass slide. A significant increase in line height and line width was observed with increasing pressure or volume flow for the two systems respectively (Supplementary Figure 3 E–H, p < 0.05). Importantly a line height of 242 μm was determined for material being extruded at 26 kPa with the pneumatic system and 226 μm was determined for material being extruded at 134.5μl/min with the PCP to avoid shifting of the material due to a too large distance of the nozzle from the print bed.

To determine the delay between start command of the gcode and the actual extrusion of material (extrusion delay), lines of 15 mm were printed at a height of 242 μm and 226 μm with the pressure system and PCP respectively. The delay was then calculated from lines printed at different delay times and data later interpolated to achieve a total line length of 15.41 mm. The additional 0.41 mm were added as the distance from the center of the nozzle at the start point to the end point was set to 15 mm and therefore an overlap of half the nozzle diameter at the start and end had to be added. Accordingly, a delay of 155 ms and 284 ms was calculated for the pressure system and the PCP respectively (Supplementary Figure 3 I, J).

#### 4) Layer Height

When printing lines next to each other, these lines fuse depending on the flow and viscoelastic behavior of the bioink and the overlap of the individual strands. Significant differences were found in layers printed with the pressure system for all three layers (layer 1: 441 ± 46 μm, layer 2: 319 ± 22 μm, layer 3: 379 ± 49 μm, p < 0.005) with an average total layer height of 380 ± 75 μm. No differences in layer height were observed in layers printed with the PCP (layer 1: 393 ± 13 μm, layer 2: 407 ± 41 μm, layer 3: 410 ± 62 μm, p > 0.85) having an average total layer height of 403 ± 35 μm (Supplementary Figure 3 K, L).

### 3.4 Comparison of the accuracy and precision of the pneumatic system to the progressive cavity pump

Cubes designed based on the results of the preparation process were printed to compare the accuracy and precision of the pneumatic system to the PCP (Figure 2 A). For the pneumatic system, a pressure of 26 kPa, a delay of 155 ms and a layer height of 380 μm were used. For the PCP, the same calibration data which were obtained in the preparation process were used, i.e. calibration was not performed again but saved values used, a volume flow of 134.5 μl/min set, a delay of 284 ms and a layer height of 403 μm used. The reduced layer height of 380 μm for the pneumatic system was chosen to ensure that the strands of subsequent layers would be in direct contact with the previous layer when deposited. As the standard deviation of the layer height of the pneumatic system was larger (±75 μm) compared to the layer height of the PCP (±35 μm) (Supplementary Figure 3K, L) using the same layer height for both systems, would risk that layers printed with the pneumatic system might not be in direct contact with the previously printed layer. Nonetheless, as the volume flow of the systems is independent of the layer height, this difference might only influence the print quality, e.g. by moving the nozzle tip through the top part of a printed layer.

The pressure system achieved an overall accuracy of 12.4 ± 4.3% and a precision of 18.99 ± 16.23 μl (batch 1: 17.39% and 12.56 μl, batch 2: 10.1% and 30.52 μl, batch 3: 9.8% and 44.96 μl) but did clog twice, once for batch 2 and once for batch 3 (Figure 2 B). When clogged, the pressure had to be increased to extrude the material clogging the nozzle or if the material would not extrude at increased pressure, the nozzle would have to be replaced. The PCP on the other hand achieved an overall accuracy of 0.3 ± 0.2% and a precision of 0.54 ± 0.45 μl (batch 1: 0.3% and 1.12 μl, batch 2: 0.2% and 0.90 μl, batch 3: 0.5% and 0.26 μl) without clogging (Figure 2 C). Despite the higher volume flow used for the pneumatic system, samples repeatedly showed a lower volume than the expected volume together with surface defects visible by small holes/missing material on the top surface of the cubes (Figure 2 B). Overall the PCP achieved a 41 times higher accuracy (p < 0.001) and 35 times higher precision (p < 0.001) than the pneumatic system.

### 3.5 Influence of the bioink composition, printing speed and nozzle diameter on the accuracy and precision of the progressive cavity pump

Further evaluating the accuracy and precision of the PCP, three different printing setups were used and evaluated with three different bioinks (Figure 3 A). Cubes printed with a nozzle of 410 μm and a feed rate of 800 mm/min, achieved an accuracy of 1.5 ± 0.7%, 1.0 ± 0.2% and 1.8 ± 0.1% and a precision of 1.53 ± 0.36 μl, 0.76 ± 0.65 μl and 0.28 ±0.27 μl for Bioink 2, 3 and 4 respectively (Figure 3 B–D). Increasing the feed rate to 1000 mm/min but keeping the 410 μm nozzle led to an accuracy of 0.4 ± 0.3%, 0.2 ± 0.2% and 0.9 ± 0.3% and a precision of 1.35 ± 1.48 μl, 0.53 ± 0.40 μl and 0.91 ± 1.09 μl for Bioink 2, 3 and 4 respectively (Figure 3 E–G). Lastly, changing the nozzle to an 840 μm nozzle and keeping the federate at 800 mm/min an accuracy of 0.9 ± 0.4%, 1.5 ± 0.6% and 0.4 ± 0.4% and a precision of 1.86 ± 1.75 μl, 1.82 ± 0.68 μl and 2.54 ± 3.38 μl were achieved for Bioink 2, 3 and 4 respectively (Figure 3 H–J).

**Figure 3:**
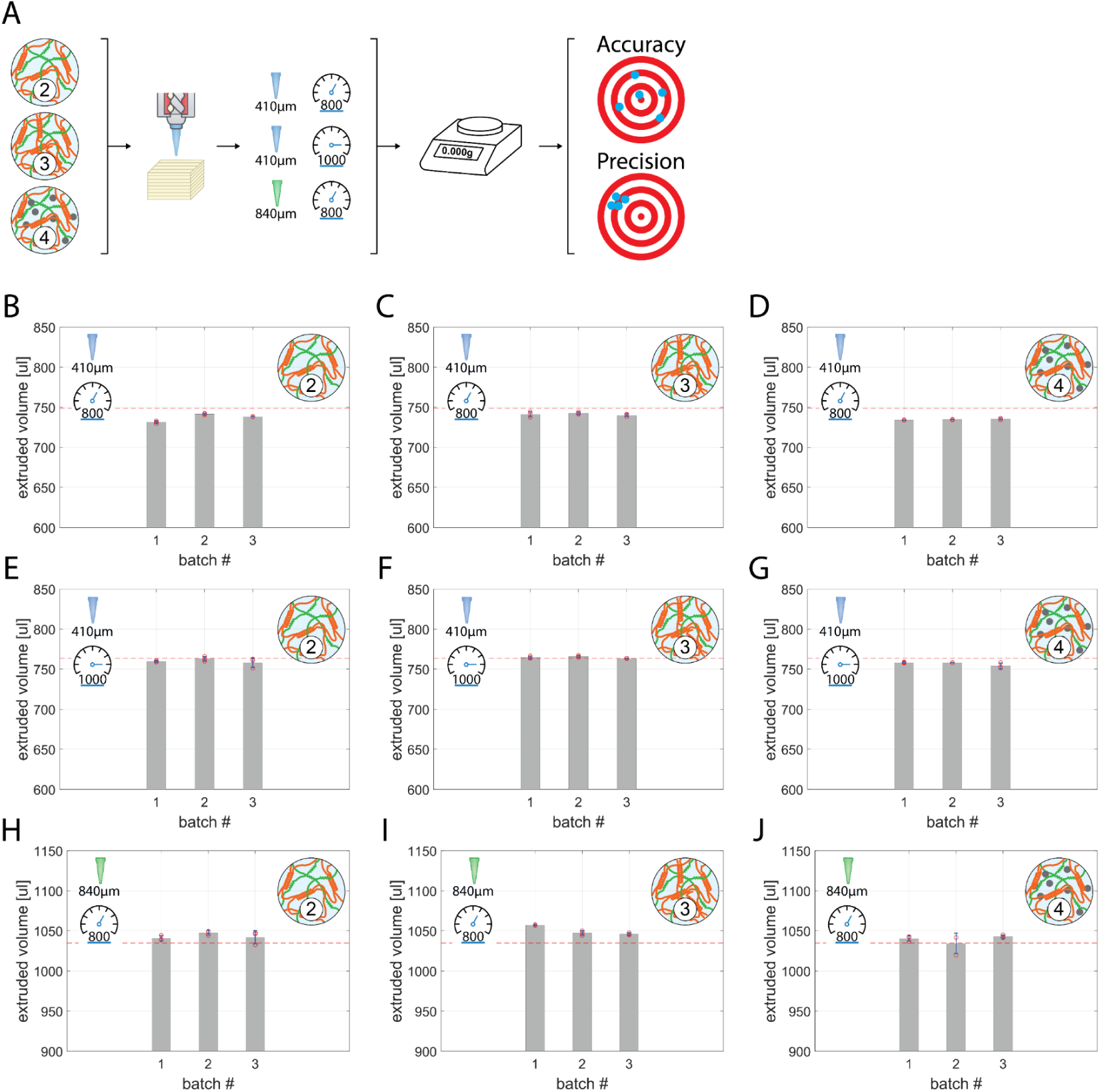
**A**, Schematic of the evaluation of accuracy and precision of the PCP. Bioinks 2–3 were used to print construcs at different printing speeds and nozzle diameter, which were afterwards weighbed and their volume compare to the calculated volume. **B**–**D**, Extruded volume of cubes printed with a 410 μm nozzle at 800 mm/min for bioink 2–4 respectively. **E**–**G**, Extruded volume of cubes printed with a 410 μm nozzle at 1000 mm/min for bioink 2–4 respectively. **H**–**J**, Extruded volume of cubes printed with a 840 μm nozzle at 800 mm/min for bioink 2–4 respectively. Red line: targeted volume.

Accuracy and precision of the PCP did not show a significant dependency on the printing parameters, i.e. nozzle size and printing speed, nor on the material used (accuracy: p= 1, precision p = 1) and no significant difference was observed compared to the accuracy and precision of the cubes printed with Bioink 1 (accuracy: p = 1, precision p = 1). Compared to the accuracy and precision of the pneumatic system, all setups used showed a significantly higher precision (p < 0.0005) and accuracy (p < 1.3e-7).

### 3.6 Viability

Viability was evaluated for casted samples and compared to the viability of samples printed with the pressure system with a 410 μm nozzle, the PCP alone and the PCP with a 410 μm nozzle (Figure 4 A).

**Figure 4:**
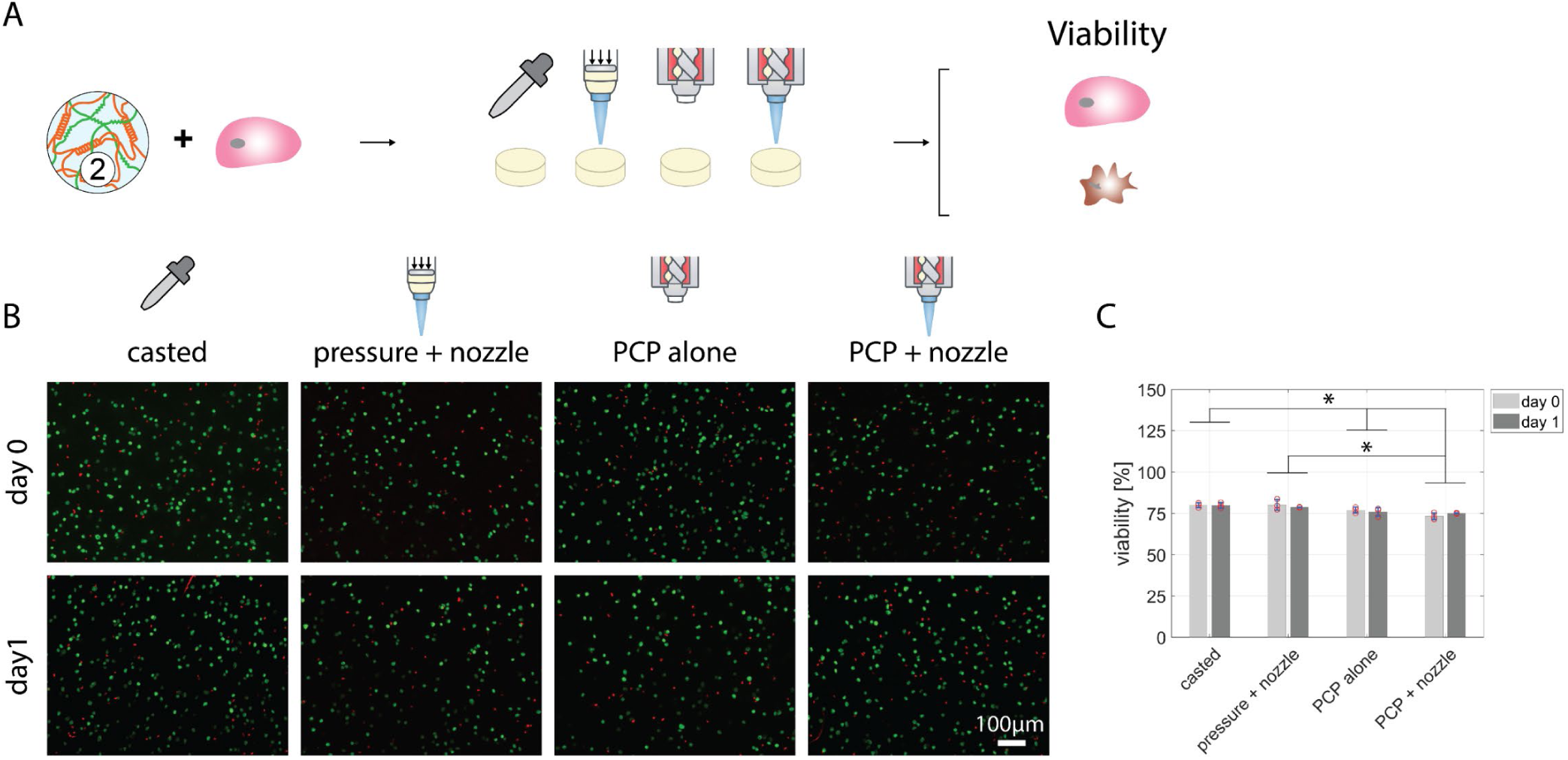
**A**, Schematic of the viability evaluation. Bioink 2 was combined with bovine chondrocytes and casted, printed with the pressure system and a 410 μm nozzle and with the PCP with and without a 410 μm nozzle. **B**, Stacked images of the viability at day 0 and 1. **C**, Viability of bovine chondrocytes at different timepoints for the different conditions.

No evidence of an interaction effect of the method and timepoints (p = 0.47) and no dependence on the chosen timepoints (p = 0.76) was detected whereas the method used significantly influenced viability (p < 0.001). Samples extruded through the PCP alone (day 0: 77 ± 2.7 %, day 1: 76 ± 2.7 %, p = 0.045) and through the PCP and a nozzle (day 0: 73 ± 2.2 %, day 1: 75 ± 2.3 %, p < 0.001) showed a significantly lower viability compared to the casted samples (day 0: 80 ± 3.1 %, day 1: 80 ± 2.0 %). Samples bioprinted with the penumatic system had a significantly higher viability (day 0: 80 ± 3.2 %, p = 1, day 1: 79 ± 1.9 %, p = 1) than samples extruded with the PCP and nozzle (p = 0.001) whereas no significant depence was detected between samples extruded through the pneumatic system and the PCP alone (p = 0.098, Figure 4 B, C).

## 4. Discussion

This study for the first time applied the PCP extrusion process to bioprinting of mammalian cells and compared the performance of a PCP to the pneumatic micro-extrusion system. Comparing accuracy and precision of the two systems, the PCP outperformed the pneumatic system in both aspects, reaching a 41 times higher accuracy and 35 times higher precision. As the pneumatic system relies heavily on a homogeneous bioink, any inhomogeneities have an influence on the material flow. Analyzing the flow rate of the pneumatic system (Supplementary Figure 3, C) it can be seen that a slight change in pressure already has a significant influence on the flow rate. Conversely, a slight change in the rheological properties of the bioink can have a significant influence on the flow rate when the pressure is kept constant. Therefore, the flow rate might be significantly altered during printing with the pneumatic system, leading to defects in the printed construct (Figure 2 B). As the PCP works based on the endless piston principle, creating an accurate and consistent flow by volumetric displacement of the created cavities, such inhomogeneities do not influence the flow rate. Once the flow rate of the PCP is determined, it can reliably and reproducibly extrude the desired amount of material. Additionally, as the PCP continuously pushes new material into the nozzle and bioinks being largely incompressible, clogging events due to inhomogeneities blocking the nozzle will be pressed out if not larger than the nozzle diameter. In contrast, once the nozzle clogs when using the pneumatic system, only an increase in pressure can ensure continuous material deposition. Such clogging events were observed twice with the pressure system, leading to failed prints. [1, 9]

The ease of use of the PCP was further evaluated by excluding the entire printing preparation process and simply calibrating the material flow for the different bioinks. Slightly larger deviations in the accuracy were observed compared to constructs printed after performing the printing assessment. These deviations could potentially be related to imprecisions in performing the calibration of the PCP or to problems related to the layer integration. If e.g. during the calibration, the weight of the extruded material is underestimated, the PCP will later extrude too much material due to a higher flow rate assumed for the material. Regarding the layer integration, the bioink flow after extrusion influences the layer height of the construct which does not reach the exact layer height of 410/840 μm. When the nozzle retracts, some of the material might not be integrated into the construct, and therefore pulled away together with the nozzle. Nonetheless, the accuracy and precision achieved with the PCP without performing the printing preparation process still outperformed the accuracy and precision achieved with the pressure system with performing the printing preparation process.

No difference in viability between casted samples and samples printed with the pressure system was observed. Previous studies showed a dependency of cell viability on the shear stresses experienced by cells during extrusion. [26] These shear stresses are in turn dependent on the material and each cell type reacts differently to different levels of shear stresses. Therefore, the reason we did not observe a drop in viability after extrusion might be explained by the shear thinning behavior of the material, reducing viscosity enough to drop the shear stresses during extrusion low enough to fall below a threshold under which bovine chondrocytes are not damaged. As we observed a drop in viability before printing (casted samples), the mixing process of the cells with the bioink appears to influence viability more significantly than the extrusion process itself. Cells extruded through the PCP alone experienced a drop in viability of 3% compared to casted samples and samples printed with the pneumatic system. As PCPs function via a pushing–and–suction action and therefore exhibit low shear rates on the material, shear stresses experienced by the cells alone might not explain the drop in viability. Additionally to the shear stresses, one has to take the seal-lines into account where stator and rotor meet to form a tight junction. At this junction cells trapped between stator and rotor might be damaged, thereby contributing to the drop in viability. Lastly, the drop in viability of 7% after extrusion through the PCP with a 410 μm nozzle might be a combined effect of the damage experienced by the cells when travelling through the PCP and then being pushed through the nozzle in which the cells experience shear stresses. Cells, which are damaged by extrusion through the PCP but would be able to recover might be terminally damaged by the shear stresses. The extrusion through the PCP might alter the shear stress threshold cells can endure, therefore damaging more cells when subsequently when extruded through a printing nozzle. To confirm these assumptions, simulations of the flow within the PCP would be necessary together with exposing the cells to controlled shear stress levels and analyzing the influence of these shear stresses on their viability.

Summarizing, progressive cavity pumps offer a valuable tool to significantly improve precision and accuracy for micro-extrusion bioprinting and therefore have the potential to replace the traditionally used pneumatic extrusion process. Their capability to extrude and maintain a precisely determined flow rate combined with knowledge about the rheological parameters of the bioink and the physical properties of the bioprinting process – acceleration and jerk of the bioprinter – can be used to optimize the printing process, thus reducing variability between and avoiding defects in printed constructs. Especially the translational aspect of bioprinting and the standardization process of bioprinted constructs would benefit from such improvements.

Future work will rely on the redesign of PCPs specifically for bioprinting to improve cell viability while maintaining their reliability and reproducibility. As materials used for bioprinting are costly, a reduction in dead volume of these pumps will be essential. From a translational perspective, all materials in contact with cellular material should either be easily cleanable or single-use. Lastly, with such an improved system, a comparison between PCPs and piston micro-extrusion to evaluate their benefits and shortcomings would further help researcher to chose the best system for their bioprinting process.

## Acknowledgements

This work was supported by the Swiss National Science Foundation (CRSII5_173868 to MZW). The authors would like to thank Annemarie Brandstetter and Raphael Lichtenecker from ViscoTec GmbH for the fruitful discussions, Dr. Nicolas Broguière for help with the rheology and Dr. Emma Cavalli for help with the cell culture.

## Supplementary Figures

**Supplementary Figure 1.**
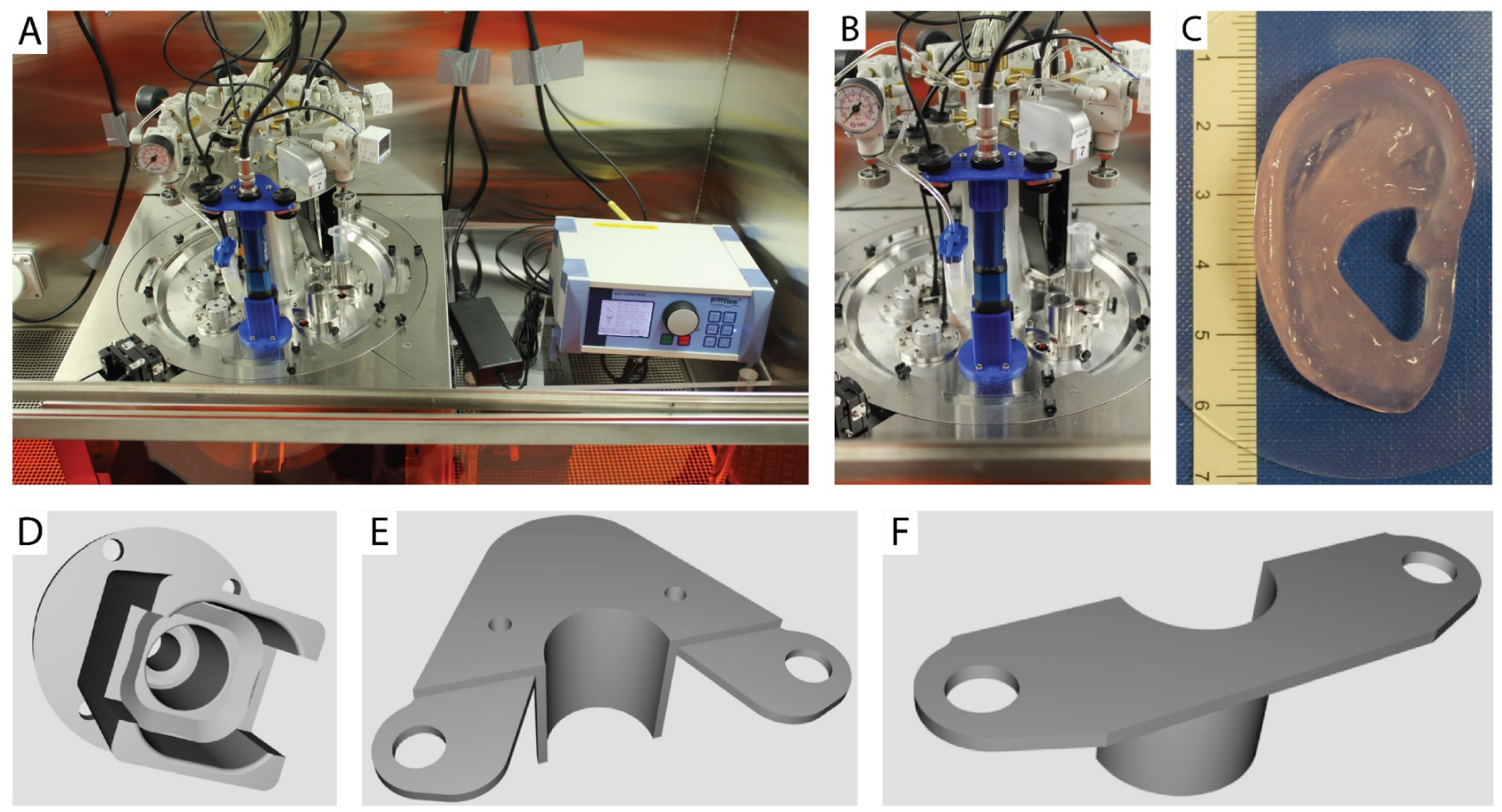
**A**, PCP and control unit installed in the Biofactory bioprinter. **B**, PCP alone mounted with the 3D printed parts on the tool changer of the Biofactory. **C**, Bioprinted ear printed with the PCP. **D**, Bottom part of the mount, holding the PCP precisely in position. **E-F**, Top parts of the mount which can be closed by screws to secure the top part of the PCP.

**Supplementary Figure 2:**
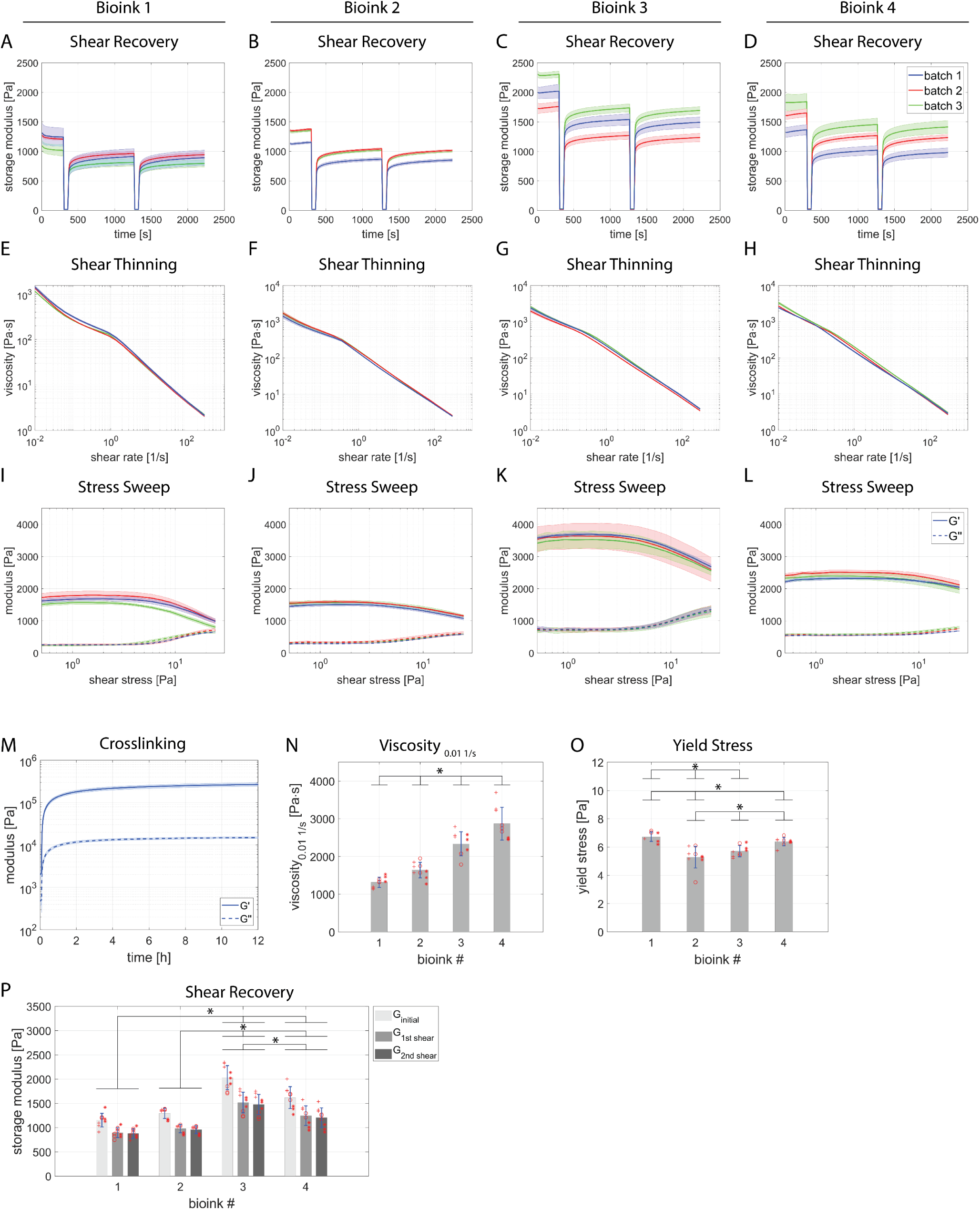
**A–D**, Shear recovery behaviour, **E–H**, Shear thinning behaviour, **I–L**, Stress sweep of the respective bioink variations. **M**, Crosslinking behaviour of bioink 1. **N**, Viscosity analysis of the different bioink variations at 0.01 1/s. **O**, Yield stress calculated from stress amplitude sweep tests. **P**, Storage moduli before shear and after the individual shear events. +: batch 1, o: batch 2, *: batch 3.

**Supplementary Figure 3:**
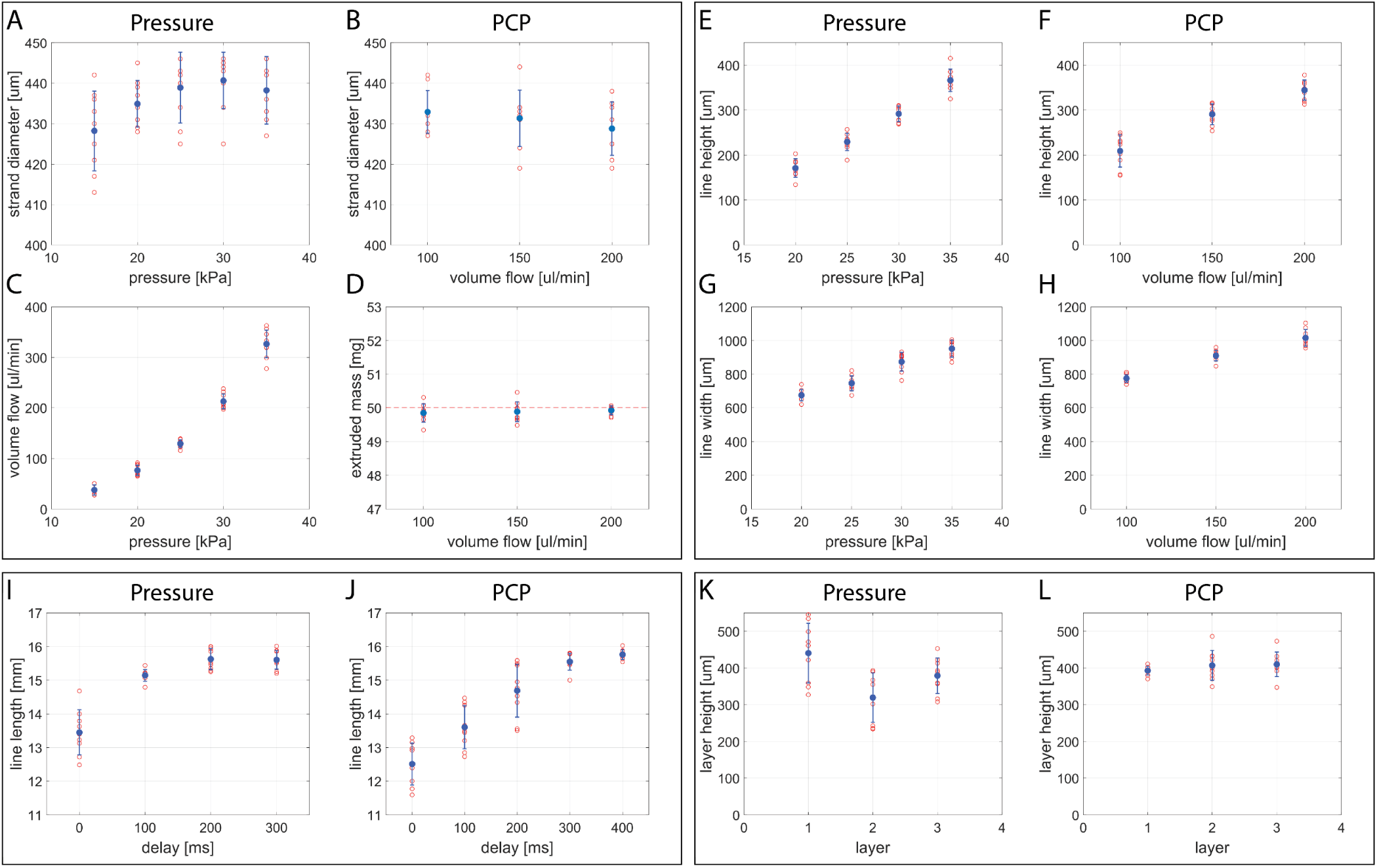
**A–B**, Strand diameter measured after extrusion through a 410 μm nozzle for the pressure system and the PCP respectively. **C**, Volume flow of the bioink at different pressures. **D**, Extruded mass after extruding a targeted 50 mg with the PCP. **E–F**, Line height of lines printed with the pressure system and the PCP respectively. **G–H**, Line width of lines printed with the pressure system and the PCP respectively. **I–J**, Line length of lines printed with different start delay times. **K–L**, Layer height of layers printed on top of each other.

**Supplementary Figure 4:**
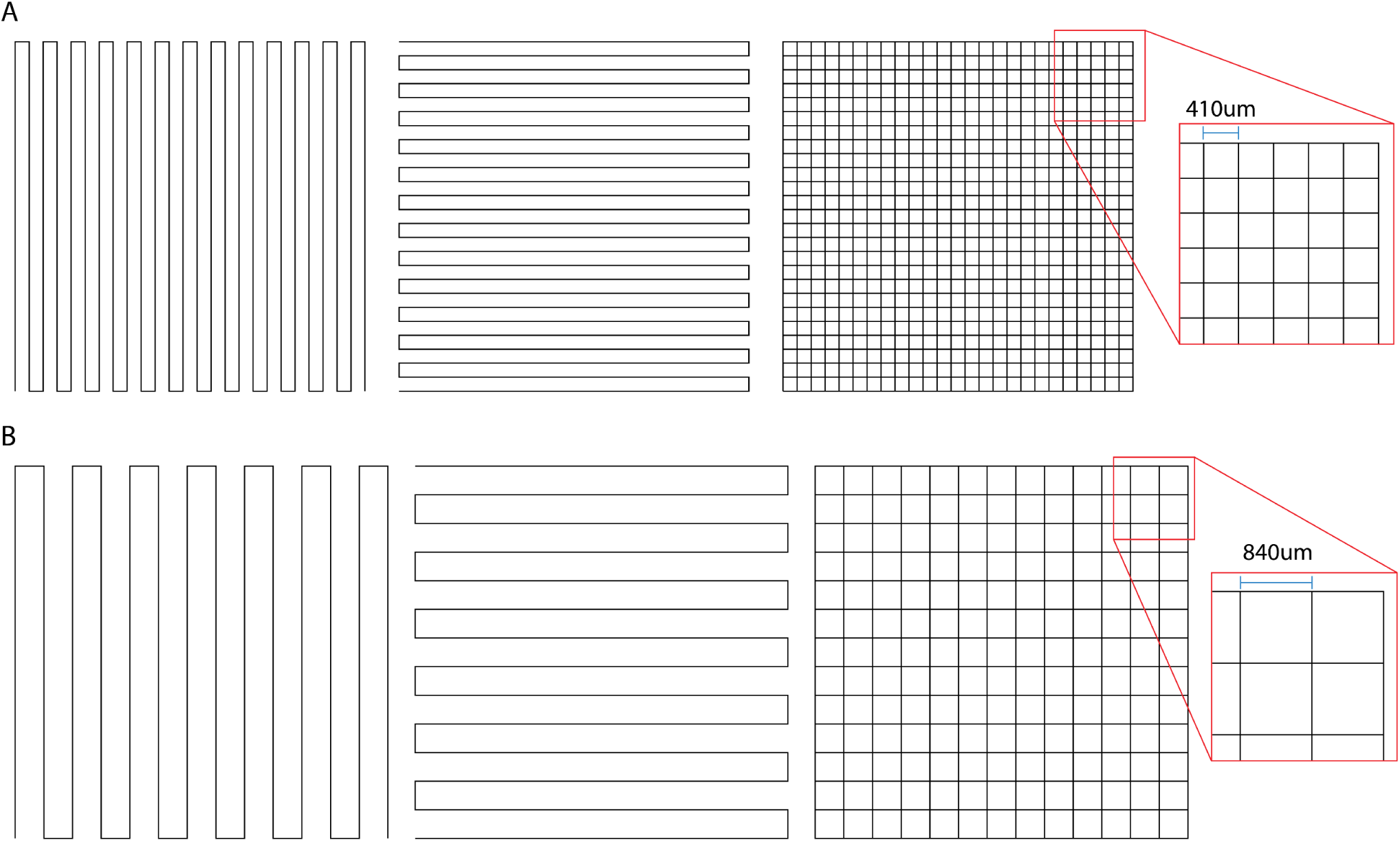
**A**, Printing path of cubes printed with a 410 μm nozzle. **B**, Printing path of cubes printed with an 840 μm nozzle.

